# The expanding truffle environment: a study of the microbial dynamics in old and new *Tuber magnatum* Picco habitat

**DOI:** 10.1101/2024.10.07.617059

**Authors:** M. Rondolini, M. Zotti, G. Bragato, L. Baciarelli Falini, L. Reale, D. Donnini

## Abstract

Truffles are valuable underground mushrooms with significant economic importance. In recent years, their cultivation has achieved satisfactory results, but not for all species. The harvesting of white truffles (*Tuber magnatum* Picco) is still dependent on natural production, which is at risk due to various issues, such as improper forest management. A useful practice to protect natural resources is to promote the expansion of productive forests. In this study, we investigate the dynamics of the microbiome in an old and new truffle forest using an amplicon sequencing approach of the fungal ITS region and the prokaryotic 16S rRNA gene. We will monitor the soil biological community’s development to compare differences and similarities between the primary productive forest and the expanding area over a two-year sampling period. In particular, we observed the colonization of vacant ecological niches by certain fungi, such as those belonging to the genus *Mortierella*. Additionally, we examined the competitive interactions between saprotrophs and ectomycorrhizal fungi (ECM). In both study areas, the bacterial community was dominated by Pseudomonadota, Planctomycetota, and Actinomycetota. The behavior of the *Tuber* genus differed significantly from other ECMs and displayed positive correlations with bacterial taxa such as *Ktedonobacter, Zavarzinella*, and *Sphingomonas*. The present work provides an initial overview of expanding white truffle habitats. Further, more specific research is needed to explore potential connections between individual *taxa*.

## Introduction

The white truffle *Tuber magnatum* Picco is a hypogeous mycorrhizal fungus. Life cycle of the fungus rely on the formation of symbiotic relationship with plants in form of ectomycorrhizas [1]. Few plants partner are able to carry the mutualistic symbiosis and depend by coevolution and niche sharing [2]. However, the whole biological cycle and reproduction of truffle depends by many other factors such as soil characteristics and climate [3]. In particular, the interplay between environmental variables as moisture, temperature, vegetation cover, soil structure and composition, and microbial component [4] determines the optimal development of ascocarps.

The truffle holds significant economic value [1], leading to potential development opportunities in rural areas [5]. White truffles thrive in environments where other crops struggle, such as valley floors. This makes them a valuable resource for preserving local communities and attracting young people to rural life. Therefore, it is crucial to study and conserve these environments, which are highly vulnerable to issues related to climate change and land abandonment.

Although the production of mycorrhized plants with white truffles is challenging, it is possible to find good quality plants in Italy and France. In spite of this, its cultivation has not yet yielded satisfactory results, unlike other truffles, so there is a growing interest in studying and conserving it in the natural environment [6,7].

Studies have been conducted on management techniques and ecological requirements [8]. While in recent years, attention has been focused on the interaction with other soil organisms, especially bacteria, which have been defined as a third partner in the symbiosis between the fungus and the plant root [9]. Numerous studies have been conducted in the identification of bacterial communities associated with truffle ascocarps [10,11]. This discovery has highlighted their significant role in producing aromatic compounds [12,13] and their involvement in nutrient cycling [14].

The advent of next generation sequencing (NGS) techniques allows to investigate the microbiome associated to truffle producing soil and ascocarps [15]. The descriptions present in literature offers valuable insights into the possible relationship between soil biological communities and truffles. However, there is a lack of comparative descriptions of the associated microbiome during the development of producing habitats. Woodland dynamics as young, nonproductive, or recently productive truffle beds may constitute an important reservoir of helpful information for *T. magnatum* sustainable exploitation as it constitutes a valuable ecosystem service due to the use of the fungus in gourmet cuisine. Normally the natural producing environments are those closest to new truffle plantations, realized in proximity with the attempt to find the ideal conditions for the development of a new truffle bed.

In our research, we have concentrated on the ecotone, which is the transitional area of the forest. This is where the young seedlings of symbiotic trees create an optimal environment for the mycelial network to grow and, within a few years, gaining the ability to produce ascocarps. We described the soil biological communities in this transitional environment between the productive forest and the agricultural field, with the aim of: i) identify the most representative *taxa* of the truffle forest and the ecotone while comparing their differences; ii) observe community changes over time: can it be assumed that the primary forest is described by a more stable microbiota than the dynamics of the secondary expansion area?

## Materials and Methods

### Study Area

This study was conducted in San Giovanni d’Asso, a location in southern Tuscany, Italy (43°9’20”52 N 11°35’27”24 E). The area is famous for its unique hillside landscapes, which are characterized by gullies: forms of slope erosion that suggest a discrete sandy component in the soil. The valleys at the base of these hills collect eroded sediments and rainfall from the slopes, defining an environment with deep, porous soils, perfect characteristics for developing white truffles [16]. The area subjected to this study was located in one of these valley bottoms, where the owner has been managing the forest for years to conserve and improve the truffle resource and where he is promoting its natural expansion bordering an abandoned cultivated field. The actions implemented are to cut and reduce the vegetation around poplar seedlings (*Populus canescens* (Aiton) Sm.), the main symbiont plant in the area, and provide water if necessary, promoting their growth and development. As already mentioned, the soil is mainly composed of sand (45%) and silt (41%), and a small percentage of clay (14%). Other soil characteristics are reported in Figure 1.

**Figure 1.**
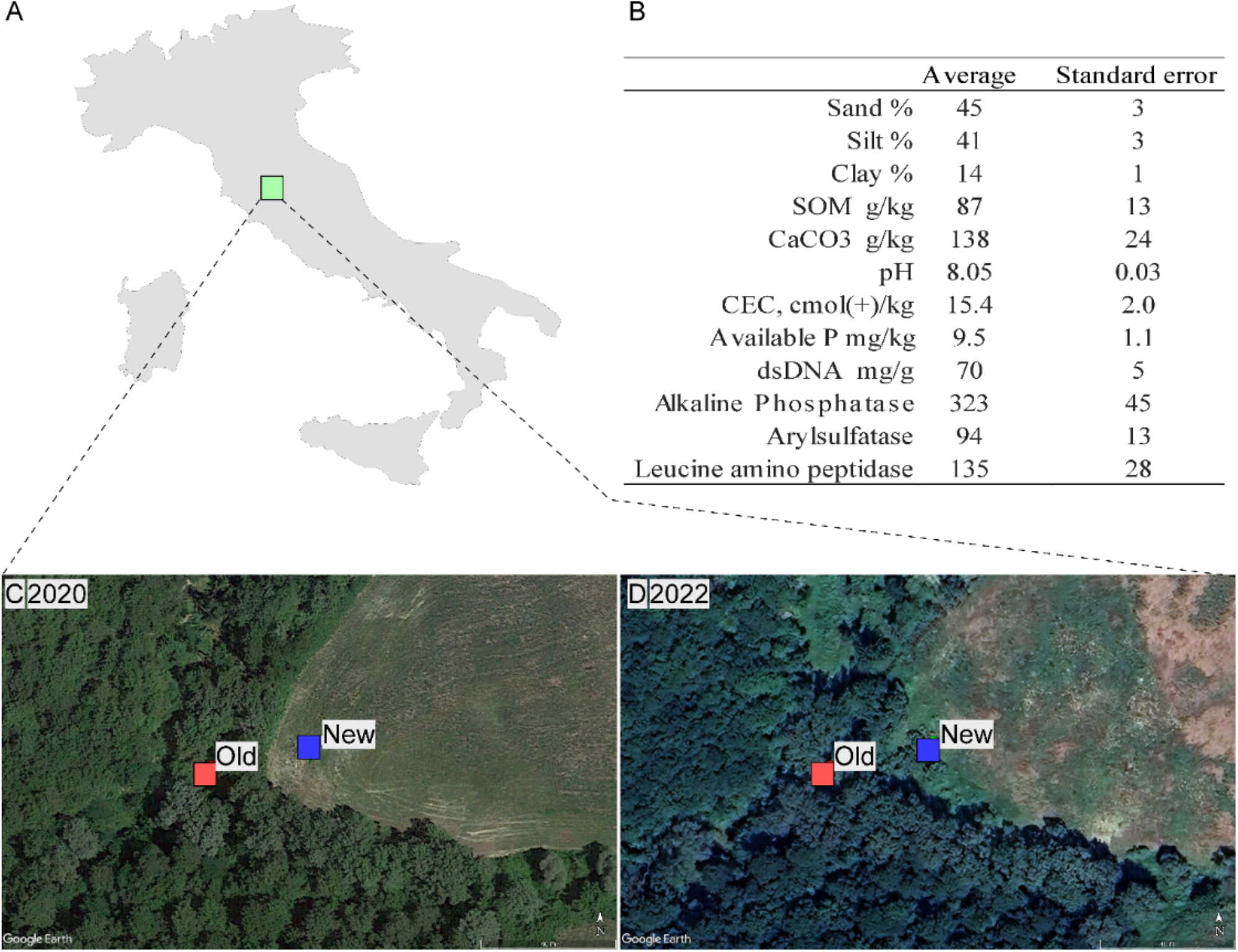
A) Location of experimental site in Italy (43°9’20”52 N 11°35’27”24 E); B) Description of soil characteristics of the study site; satellite images of the study sites in 2020 C and in 2022 D with respective Old (red squares) and New (blue square) sampling point.

### Experimental design, soil sampling and DNA extraction

With the idea of describing the microbial succession during different years of expansion of a naturally producing with truffle area, we designed the experiment as a comparison between producing woodland (Old) and the ecotonal area with expanding renewal of potential symbiont (New). In detail, we collected soil samples in autumn 2020 and 2022: 6 soil samples in the expanding area, which we will refer to as ‘New’, and another 6 in the adjacent truffle forest, which we will refer to as ‘Old’. Core samples were collected using a 16mm diameter PVC tube that had been previously sterilized. The tube was hammered into the soil to a depth of 20 cm after removing the litter layer. A total of 24 samples were collected in two sites and two seasons. After collection, the cores were transported in a portable refrigerator at 4 °C using ice packs until their storage at - 80 °C. Before molecular analysis, the samples were freeze-dried and then pounded. Samples were sieved to avoid stones and root debris with a mash size of 500 µ. The whole procedure was carried out taking care to disinfect each instrument before using and processing another sample. DNA was extracted from about 0.40 gr of soil per sample using Qiagen, Hilden, Germany, DNeasy PowerSoil Kit (Cat No./ID: 12888-100) following the manufacturer’s instructions. The DNA was eluted with 50 μL of distilled water and stored at -20°C until further analysis.

### Bioinformatic analyses

PCR analyses were conducted by Sequentia Biotech SL (Barcelona, Spain), including library preparations, Illumina MiSeq sequencing of the bacterial 16S rRNA genes and fungal ITS region, and bioinformatic analysis.

The primers used for amplification of the 16S region are 341F 5’-CCTACGGGNGGCWGCAG-3’ and 805R 5’-GACTACHVGGGTATCTAATCC-3 and for the ITS region: ITS1 5’-TCCGTAGGTGAACCTGCGG-3’ and ITS2 5’-GCTGCGTTCTTCATCGATGC -3’. Library generation was carried out following the protocol recommended by the kit manufacturer (Illumina). The raw data were subjected to a quality check using the software BBDuk. The trimming was done removing the low-quality parts of the data while keeping intact the longest high-quality part of the data. The minimum read length required for the analysis was set at 35 bp, with a quality score of 25 to ensure high-quality and reliable results. The software GAIA (version 2.02, Sequentia Biotech, Spain) was used to analyze the taxonomic profiling of the samples. The process involves aligning each set of reads with a reference database to extract the best alignments for accurate comparison, and then a Lowest Common Ancestor (LCA) algorithm is applied to identify the best alignments. The analysis returned a table of OTUs for each taxonomic level. Raw sequence reads have been submitted to the Sequence Read Archive linked to the bio-project number PRJNA1167916 in the National Center of Biotechnology Information (https://www.ncbi.nlm.nih.gov/bioproject/).

### Statistical Methods

Alpha diversity indices were calculated using a *vegan* package [17] with the “specnumber” function for the richness and “diversity” function for Shannon’s index calculation. Also, Number of reads for each sample were used as indicator of alpha diversity. Significative changes in Species richness, reads number and Shannon index were tested by means of two-way analysis of variance (two-way ANOVA).

Beta diversity between the samples was calculated with the Bray-Curtis dissimilarity index again using the *vegan* package with the “vegdist” function. For both bacterial and fungal communities, Bray-Curtis dissimilarity matrix were calculated and multidimensional ordination of samples visualized by Non-metric MultiDimensional Scaling (NMDS). Ordinations were performed with the “metaMDS”, Permutational analysis of variance (PERMANOVA) was used to quantify the impact of variables on dissimilarity between communities with “adonis2” function. To observe the taxa-specific variations in bacterial and fungal communities, heatmaps were built comparing New and Old sites across different seasons for both bacterial and fungal communities restricting the analysis on the 70 more representative *taxa*. To identify microbial groups according to their abundance in experimental design, variables were ordered using hierarchical clustering based on Index of association. Furtherly, volcan plot were built to isolate the main statistically significant *taxa* producing samples differentiations. Metrics for volcan plots were calculated as fold changes across years for NEW and OLD sites and significance calculated by means of t-test of homoscedastic data.

The most abundant fungi in the different areas, classified at the genus level, were annotated with their trophic function via the FUNGuild database using the ‘funguild_assign’ function in the *FUNGuildR* package [18]. FUNGuild data, frequence of *Tuber* genus, and most representative bacteria obtained by volcan plot were checked for correlation by means of Spearman rank values with the aims to understand microbiological dynamic and possible relationships in Old and New forested soils. The resulting correlation maps were ordinated for fungal trophic groups and bacteria by means of hierarchical clustering based on Euclidean distance metrics. Statistical analyses were carried out in the R environment (R Core Team 2020) and using the Primer 7 software (PRIMER-E Ltd, Plymouth; UK).

## Results

18.431.100 OTU and 10.928.616 OTU were obtained respectively for Bacteria and Eukaryota domains. Taxonomic analysis at the phylum level showed minimal differences in Bacterial community, while more marked for Fungi (Fig 2).

**Figure 2.**
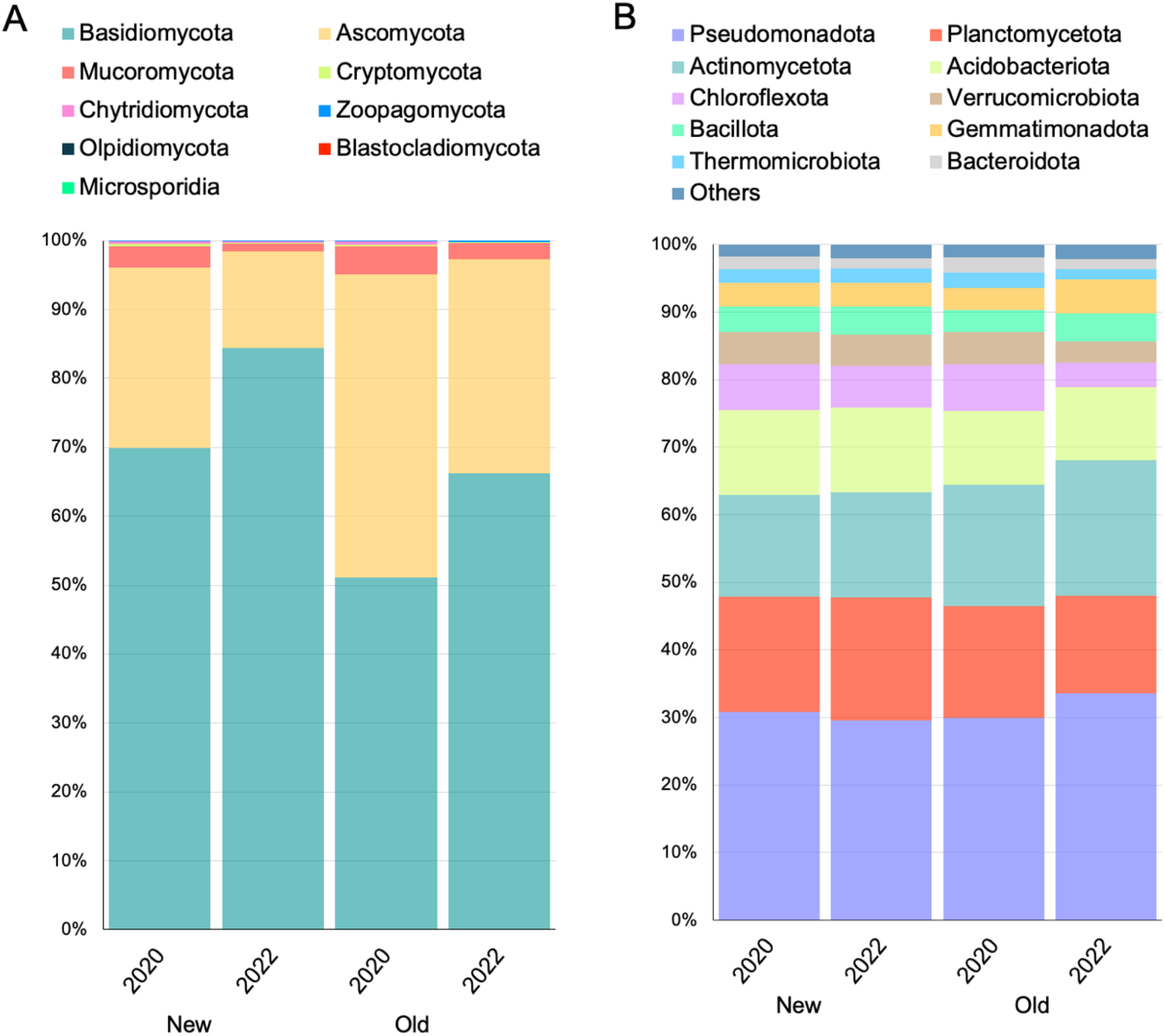
Stacked bar chart of the identified OTUs at the phylum level. A) The percentage of detected fungal phyla; B) The percentage of detected prokaryotic phyla.

For the fungal community, members of Basidiomycota are the most representative *taxa*. Specifically in New site the presence of Basidiomycota is more abundant in expanding area compared to the old one (77.12% New vs 66,22% in Old). Moreover, increase of Basidiomycota frequence is observed according to year being higher in 2022 when comparing seasonality across the same experimental area. The second and the third most representative phyla were Ascomycota and Mucoromycota, respectively, those showed the same pattern of change in terms of frequence, mostly due to the years, with a sharp decline in 2022 for both New and Old areas.

Pseudomonadota, Actinomycetota, and Planctomycetota are the predominant bacterial taxa in the study areas. The former showed a differential effect according to the year and the site being particularly favored in Old area in 2022 (33,55%) while depressed in New area of the same year. Actinomycetota are disadvantaged in the expanding area (∼15% in New vs ∼19% in Old) but does not undergo great changes over the years within the same site. Instead Planctomycetota showed a distinct association with the Expanding area compared to the Old one (∼17,5% in New vs ∼15% in Old).

### Alpha and Beta diversity

The alpha diversity analysis for the fungal community showed no significant change for Species richness across years and sampling areas (F=2,51; p=0.087), however a notable decrease is observed in some sample from Old area in 2022 (Figure 3A). Inversely, Number of reads significantly decrease in Old area compared to the New ones with no effect observed for the years (F=3,25; p=0.042; Figure 3B). Shannon index values also change significantly, showing a dominance pattern marked by lower values in New and Old areas in 2022 (F=3,14; p=0,047; Figure 3C). Non-multi-dimensional scaling (NMDs) of New area showed that fungal community do not undergoes to complete diversification across years. However fungal community appear more similar between sampling points in 2022 while more dissimilar within 2020 (PERMANOVA, PER=999, p = 0.019; Figure 3D). Same ordination method for samples of Old area denotes, instead, a clear differentiation in fungal community across years (PERMANOVA with 999 perm. p = 0.002; Figure 3E).

**Figure 3.**
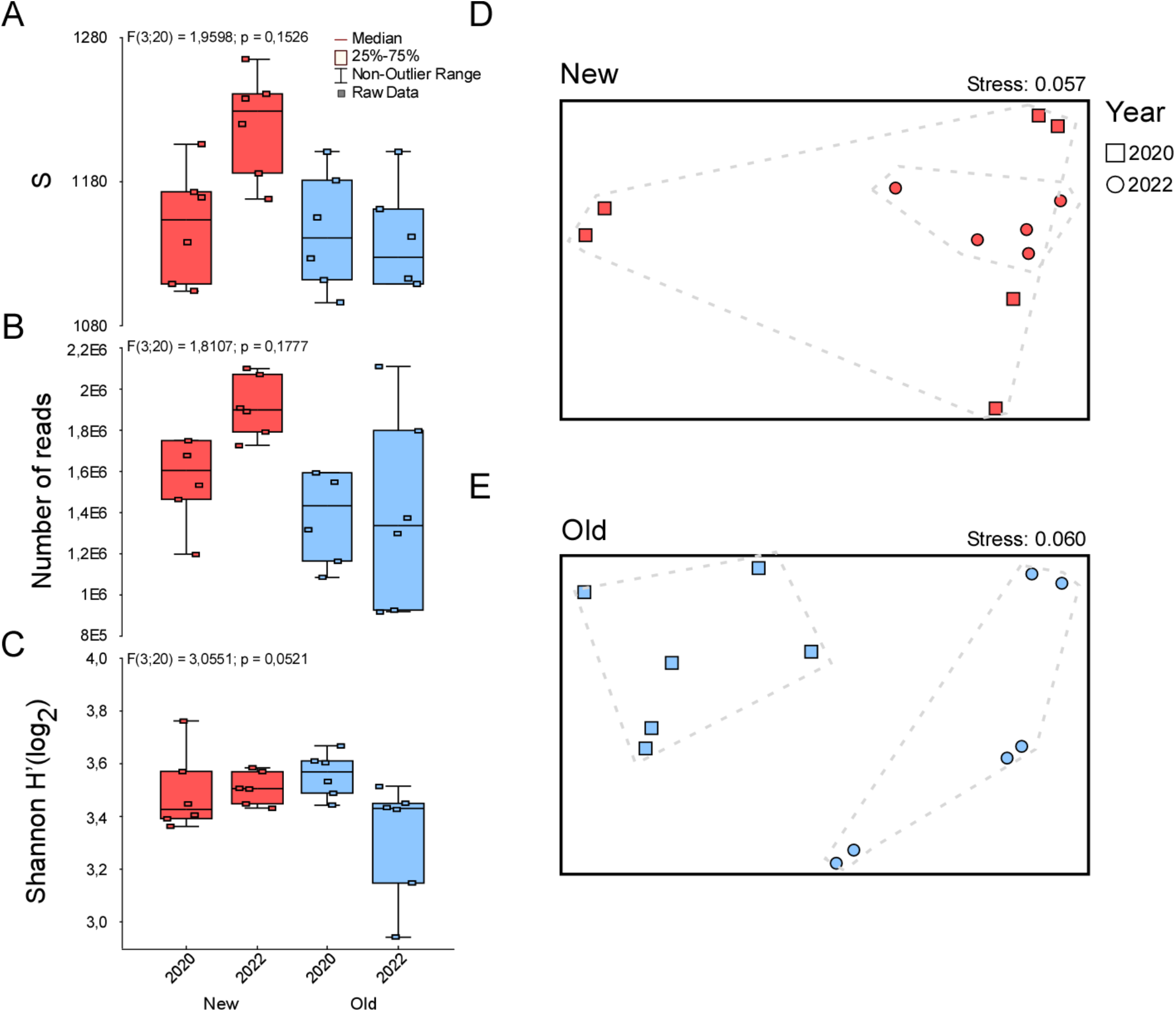
Alpha diversity boxplots showing Species richness, Number of reads, and Shannon index for the fungal community for each site in different years (A– C). Non-metric MultiDimensional Scaling (NMDS) plots and relative stress level, using Bray-Curtis dissimilarity matrices of fungal community for New site (D) and Old site (E).

For bacterial community, alpha diversity metrics shows no significant differences between samples. However, increase of bacterial richness and Number of reads is observed in the New areas of 2022 (Figure 4 A, B and C). Specularly for what observed in fungal community for both New and Old areas. In New area NMDs showed overlap of community similarities between 2020 and 2022 but with a marker clustering of 2022 sites (PERMANOVA with 999 perm. p = 0.067; Figure 4D). In Old area, as also observed for fungi, clear differentiation of bacterial community occurs between 2020 and 2022 (PERMANOVA with 999 perm. p = 0.004; Figure 4E).

**Figure 4.**
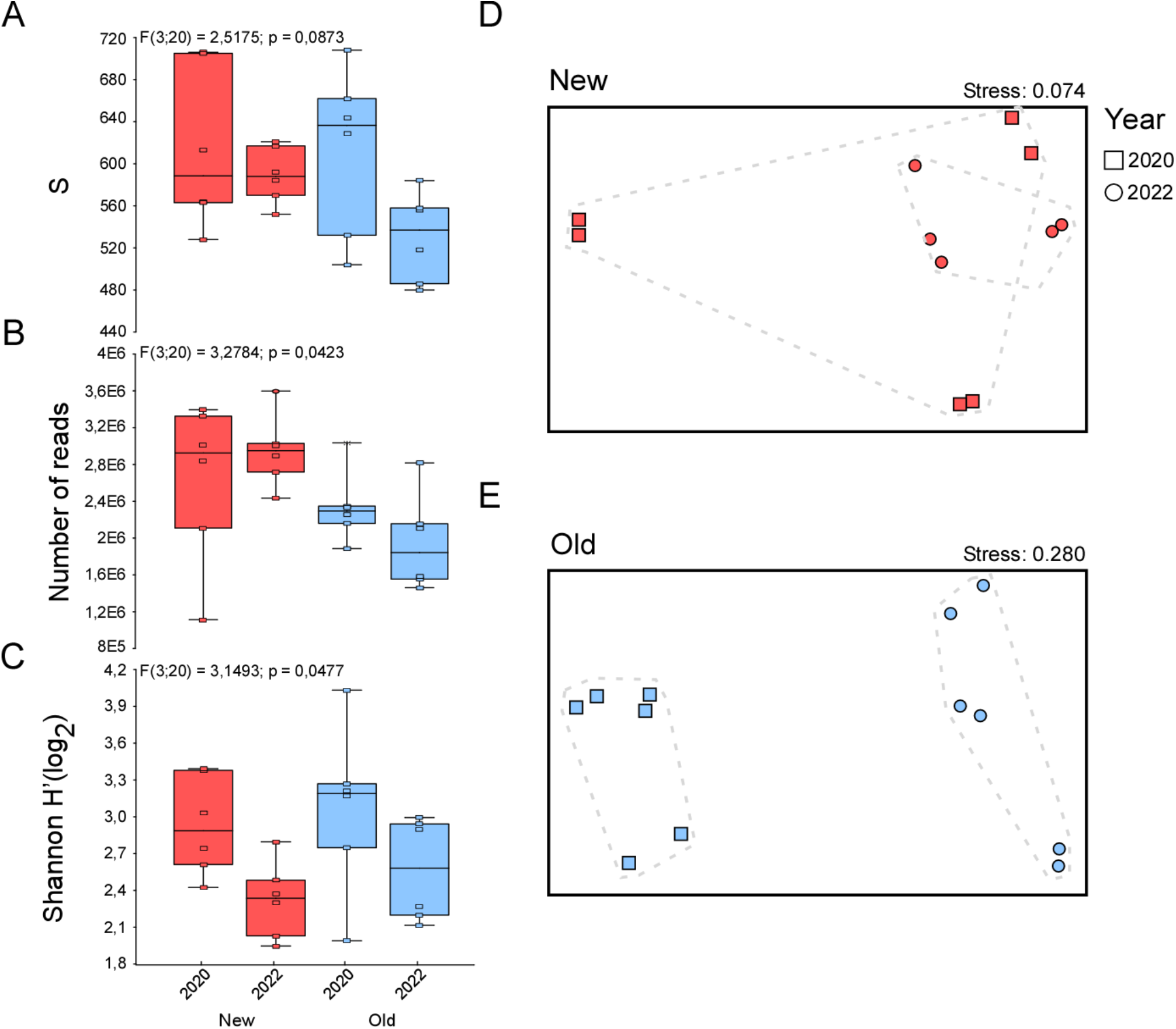
Alpha diversity boxplots showing Species richness, Number of reads, and Shannon index for the prokaryotic community for each site in different years (A– C). Non-multi-dimensional scaling (NMDs) plots and relative stress level, using Bray-Curtis dissimilarity matrices of fungal community for New site (D) and Old site (E).

### Microbiome associated to expanding truffle habitat

*Taxa* contributing to differentiation of sampling areas are showed by means of Heatmaps and volcan plots for both fungal (Figure 5) and bacterial (Figure 6) communities.

**Figure 5.**
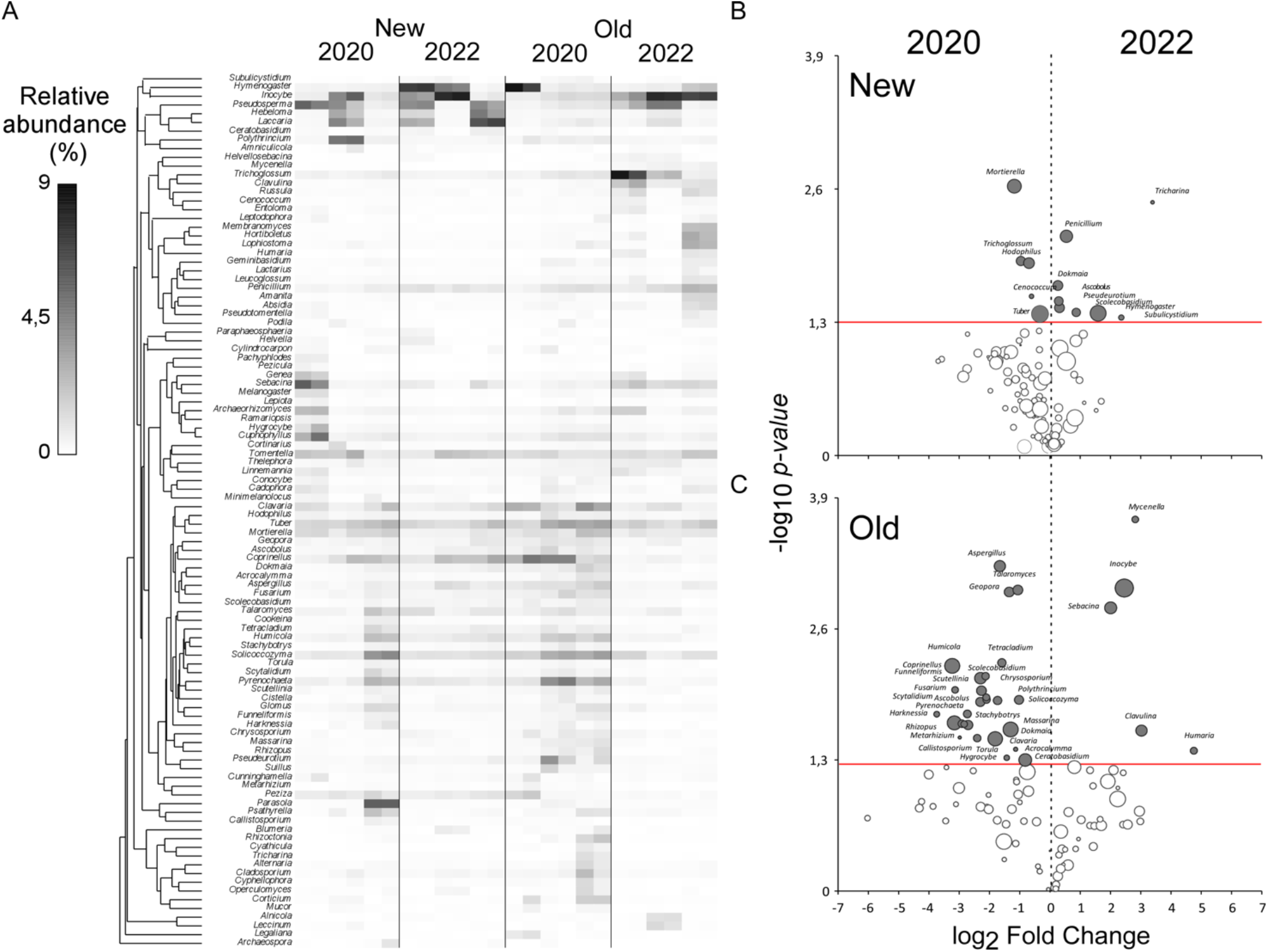
A) Heatmap showing the relative abundance of the 70 most representative fungal taxa for each site in different years. Hierarchical clustering is based on Index of association; The volcan plots demonstrate the patterns of enrichment and diminishment in the fungal community through the years in New site (B) and in Old site (C).

**Figure 6.**
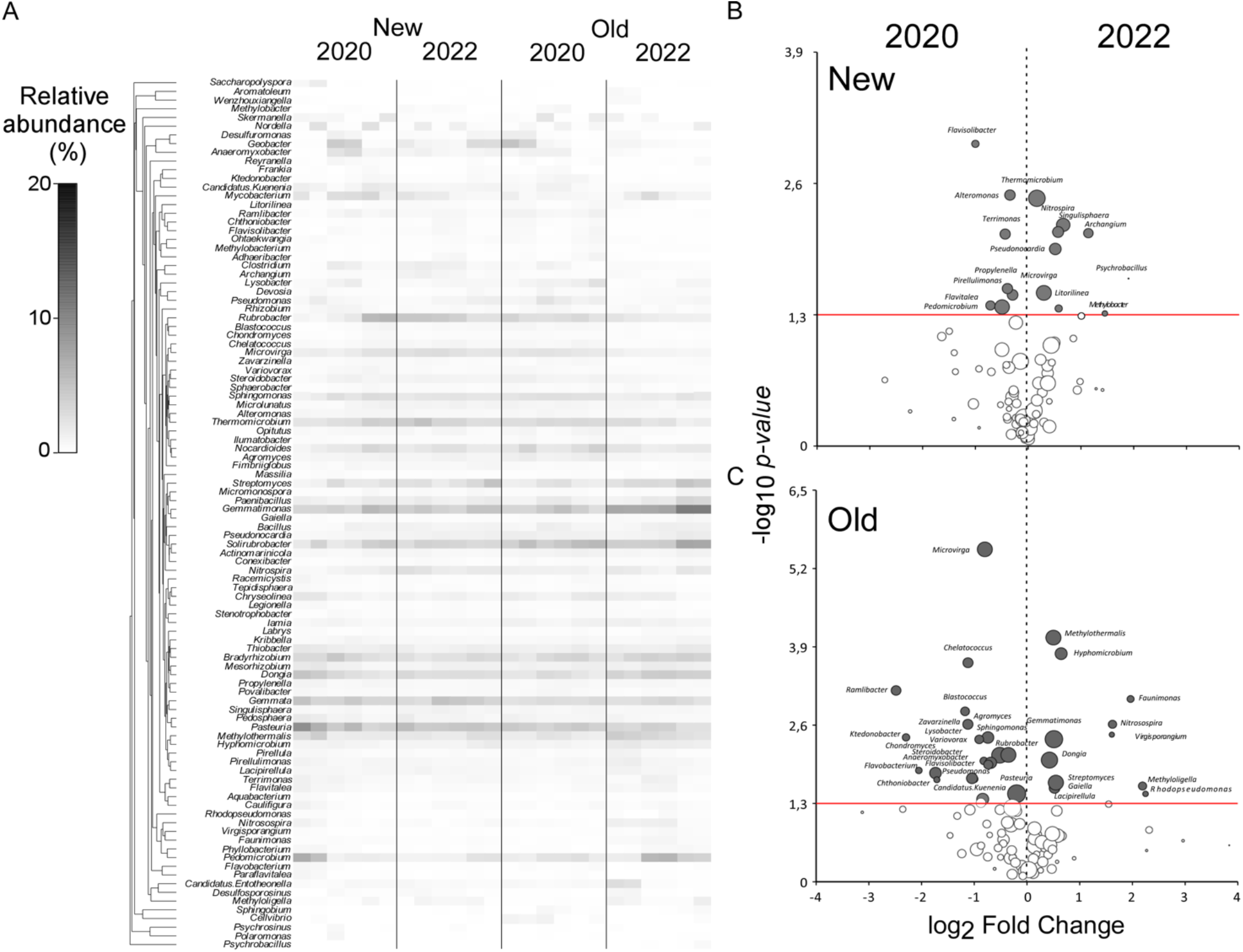
A) Heatmap showing the relative abundance of the 70 most representative prokaryotic taxa for each site in different years. Hierarchical clustering is based on Index of association; The volcan plots demonstrate the patterns of enrichment and diminishment in the prokaryotic community through the years in New site (B) and in Old site (C).

Among fungi, New areas are differentiated because of the high frequency of *taxa* belonging to *Tuber, Cenococcum, Hodophilus, Trichoglossum and Mortierella* in 2020 (Figure 5B). In 2022 those genera were significantly substituted *by Subulycistidium, Hymenogaster, Scolecobasidium, Pseudeurotium, Ascobolus, Dokmania, Penicillium* and *Tricharina*. Notable increase, despite non significative, was also the increase in 2022 of *Inocybe, Laccaria, Penicillum*, and *Coprinellus* (Figure 5A). Within Old areas 2020 were characterized by higher frequency of *Aspergillus, Talaromyces, Geopora, Humicola, Tetracladium* and *Coprinellus*. The fungi of the genus *Tuber* also showed increase in relative abundance for 2020 but not in significative way. In 2022 the community shift was mainly characterized by the higher abundance of *Inocybe, Mycenella, Sebacina, Clavulina* and *Humaria* (Figure 5C). Notably, also fungi of the genus *Pseudosperma* considerably increase in relative abundance despite not at significative level.

In New area Bacterial community across different years changed because of the high presence of *Flavisolibacter, Alteromonas, Terrimonas, Propylenella, Pirellulomonas, Flavitalea* and *Pedomicrobium* in 2020. The bacterial composition of soil shifted in 2022 toward a community mainly composed by *Thermomicrobium, Nitrospira, Singulisphaera, Archangium, Pseudonocardia, Microvirga, Litorilinea* and *Methylobacter* (Figure 6B). In Old habitat the change across different years were mainly due to high frequence of *Microvirga, Chelatococcus, Ramilibacter, Blastococcus, Abromyces Sphingomonas, Rubromyces* and *Pasteuria*. While in 2022 *Methylothermalis, Hyphomicrobium, Gemmatiomonas, Dongia* and *Streptomyces* were favored (Figure 6C).

### Ecological dynamics of mycobiome and associates bacterial consortia

Staked bar plot of FunGuild data showed different fungal community pattern when grouped according to their trophic strategy (Figure S1). The differentiation is mainly due to the different year of sampling as in 2022 dominance of Ectomycorrhizal (ECM) fungi were recorded for both New and Old areas. When comparing the year in the same habitat is observed for new site that the ECM dominance excludes many saprobic *taxa* as well as plant pathogens, Epiphytes and in minor extent animal parasites. The results of New site mirrored those observed in Old habitat, as in 2022 ECM dominance is recorded becoming disadvantageous for plant pathogen, saprotrophs, Epiphytes and others minor fungal groups.

Accordingly, when clustering fungal guild together, euclidean distance values showed that endophytes and fungi belonging to the *Tuber* genus has similar pattern in forest soil (Cluster1) While animal pathogens, saprotrophs, plant pathogens and epiphytes clustered together (Cluster 2), while other groups as animal, plant saprotrophs, Fungal parasites, Arbuscular mycorrhizal fungi and wood saprotrophs showed no strong associative patterns each other (Cluster 3). ECM groups showed complete opposite effect compared to the other fungal groups (Cluster 4) (Figure 7). When defining bacterial consortia associated to fungal trophic group is observed that Cluster 1 associated with Bacteria of the *Ktedonobacter, Zavarzinella* and *Sphingomonas* genera while Cluster 2 associate with *Blastococcus, Rubrobacter, Chondromyces, Thermomicrobium, Microvirga* and others. Cluster 3 showed no notable correlation with microbial groups, while ECM (Cluster 4) positively correlates with *Lacipirellula, Dongia, Hyphomicrobium, Singulisphaera* and *Archangium* genera.

**Figure 7.**
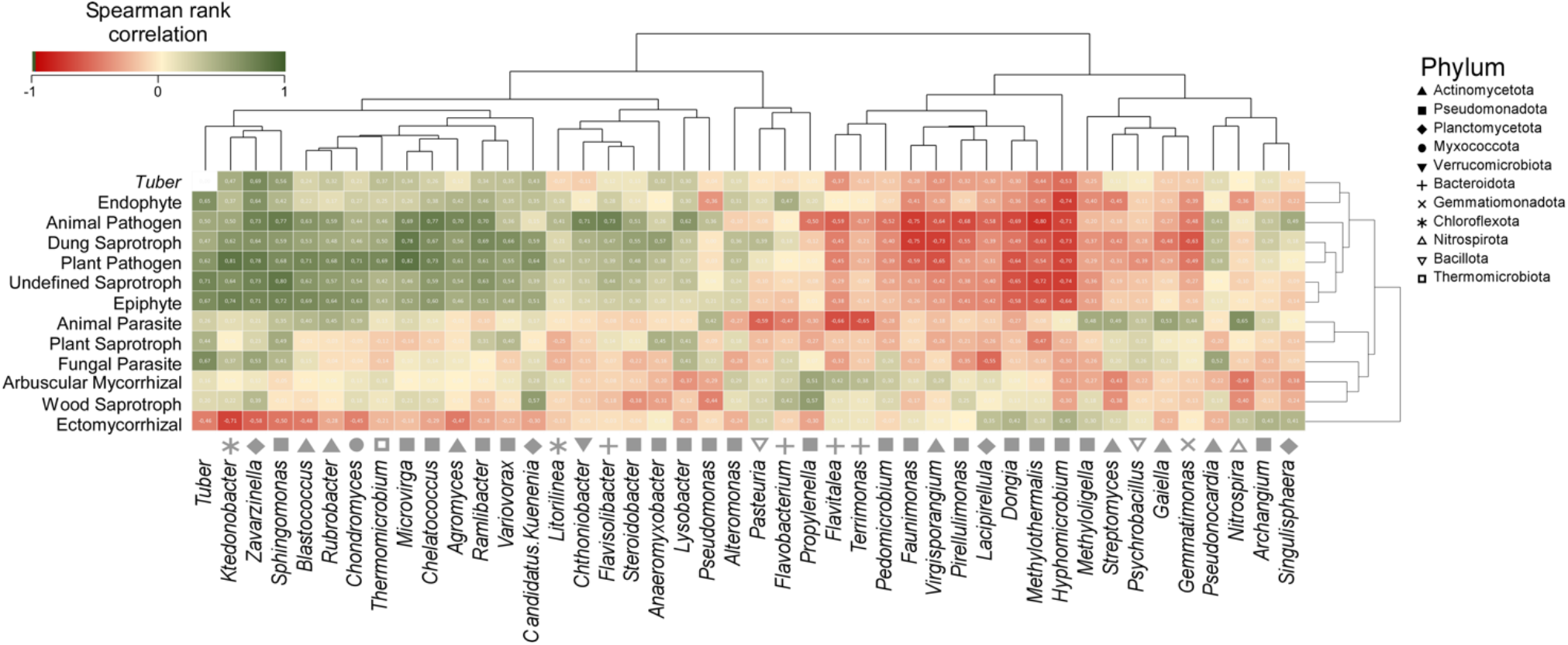
Spearman’s rank correlation maps showing the relationships between *Tuber* genus, fungal trophic groups and bacteria. Hierarchical clustering is based on Euclidean distance metrics.

## Discussion

In recent times, several studies have focused on the relationship between the environment and truffles, to understand some key associations to gain insight into the biology of the species [13,15,19-22] In dynamic environments such as truffle forests, the same biological factors, fungi and bacteria, follow dynamics dictated by seasonality and their relationship. In a past study, Lalli et al. (2015) [23] noted an affinity of *Amanita stenospora, Cortinarius aprinus, Hebeloma quercetorum* and *Hygrophorus arbustivus* var. *quercetorum* for the white truffle habitat, while *Mortierella* and *Fusarium* were found abundant in truffle soil by Mello et al. (2010) [24]. In this study, we investigated natural dynamics in truffle forests across two years of sampling and its expansion into the nearby abandoned field.

### Fungal dynamics in expanding truffle forest

The most evident results of our survey on fungal community at phylum level regard the increase in relative percentage of reads belonging to Basidiomycetes. Those vary across years, as rainfalls and temperatures during the 2022 may has been more favorable for mycelial expansion of this phylum. Accordingly, the dominance of Basidiomycota in 2022 is coupled with the overall increase in dominance within alpha diversity and decrease of evenness among fungal *taxa*, which means few *taxa* proliferating in soil environment as result of out-competition of the previous community. The effect of dominant mycelial mats by Basidiomycetes has been described in literature during process of organic matter decomposition and in grassland environment for free living soil saprotrophs [25-28]. Basidiomycetes intervene in the last successional step of organic matter decomposition and habitat colonization [29,30] as they are more prone to degrade recalcitrant molecules as Lignin, Celluloses and Phenols because of the wider enzymatic arsenal and oxidative trophic strategies [31,32]. Indeed, in this group of fungi, many are considered as combative species monopolizing resources and defending it by potential competitors [33,34].

When considering the internal variability among New and Old forest we observed that fungal diversity is higher in New area compared to the Old with net distinction in community composition due to years for the last. For the former, instead, communities in 2020 partially overlap with those of 2022 but is mainly coupled with a large dissimilarity of community within the sampling group. This indicate that in the early stage of colonization of a potential forest niche fungal communities can assume different shapes with a heterogeneous community composition [25]. This probably depends by the first arrival and establishment within the new empty niches, that is unpredictable in terms of *taxa* composition [35]. With time, and establishment of forests environment, fungal community stabilize and homogenize its composition in parallel with plant cover and species normalization [36,37]. Interestingly, the New expanding area in this study were managed by the owner favoring potential plant host for *T. magnatum* colonization. The results observed may be caused by this selective practice [36]. Remarkably, the alpha diversity indices, confirm the high variability of the New forming area and evidenced that on the ecotone fungal species richness and number of reads is tendentially higher with respect of the Old area. The last, is instead, characterized by lower variability because of higher time for fungal community establishment since environment formation. Hence, the higher diversity in New area may be due to the opportunistic colonization by many ruderal species that is a general rule in the ecology of colonization of new environment at multi-Kingdom level [27,38]. When considering the change in New area at higher taxonomic level (genus), the specific changes were not clearly visible as higher variability within the 2020 decrease the amount of *taxa* with significant frequency shift. However, fungi of the genus *Mortierella* provide a clearer signal of opportunistic colonization of empty niches, as has been described in many disturbed or not structured soil environments [39]. On the other side, in 2022, the significative expansion of *Hymenogaster* [40,41], an ECM forming fungus, conciliate with the observation made at phylum level regarding the dominance of the Basidiomycota phylum. Notably, in 2020 high frequence of reads belonging to *Tuber spp*. were found indicating a favorable season and suggesting a pioneer colonization of new forming environments characteristic of the genus [42,43]. In Old environment the community stabilization during the whole life cycle of the habitat results in a clearer representation of the community shift during the years. In 2020 the community was composed by a high variety of fungal *taxa* with disparate trophic guild that are promptly substituted by a poorer community that is, instead, dominated by the mycelium of ECM species mostly *Inocybe, Humaria, Sebacina*, and *Clavulina* genera [40,41,44]. Those observations at genus scale are also confirmed by an analysis at guild level that evidence an important enrichment in soil of ECM mycelium across years. The ecological implication of the finding may explain the pattern, in stable ECM woodland environments the presence of ECM symbiothrophs became dominant. When environmental conditions became favorable for ECM mycelial exploration of soil space exclusion of many saprobic species in a process called Gadgil effect take place [45,46]. Competition between the two fungal guilds for organic nutrients and other soil resources is believed to result in the suppression of the decomposition rate of organic matter (SOM), favoring ECM lifestyle. Firstly, it is important to note that the samples were collected at a depth of 20 cm, which could potentially distinguish between the ecological niches of the two groups. Saprophytic fungi are typically found in the superficial part, where there is more SOM, while ECMs are generally located at greater depths, except for a few species [47,48]. This may be due to an increase in SOM, which decreased in 2022 when the landowner cut the vegetation. The area is managed by periodically cutting excess shrub vegetation to promote continuous forest rejuvenation. According to this principle, the ecotone has also undergone selective cutting, favoring the growth of young symbiotic plants. The removal of most of the plant residues may result in less SOM, thus favoring the presence of ECM fungi.

### Bacterial dynamics in expanding truffle forest

In the prokaryotic community, we observed repetition and certain level of stability in the dominant *taxa*, which *Pseudomonadota, Planctomycetota*, and *Actinomycetota* represented. However, bacterial dynamyc in the survey design partially mirrored what observed from the analysis of fungal community. Particularly from beta diversity. This lead to think that the few changes observed are linked to the expansion of mycelium of certain *taxa* of fungal guild in soil. In recent years the attention towards the relationships between bacterial *taxa* and fungal mycelium increased, also because facilitated by the advent of NGS sequencing [49,50]. In detail, the changes observed in our surveys indicate that *Pseudomonadota* (Proteobacteria), is the bacterial phylum that predominating in the study areas, and particularly favored in Old area in 2022. Those *taxa* are normally associated to the release of nutrients in the environment as many of those species are copiotrophics with strong dependence on Nitrogen budget [26,51]. It may be argued that during the period of stronger mycelial activity (2022) the decomposition of organic matter acted by dominant ECM community may release nutrients in simplified way. Indeed, in the area with the higher frequence of *Pseudomonadota*, members of *Hyphomycrobiales* thrive including bacterial belonging to *Methilobacter, Dongia, Hyphomicrobium, Archangium* and *Methilotermalis* genera. Interestingly, those *taxa* showed positive correlation with the abundance of ECM species in the soil. On the other side, the high amount of fungal mycelium and its senescent portion may release high level of nutrients in soil environment [27,51]. The relationship with the dominant decomposition ability of ECM species found further suggestion in the presence of Bacterial genera specialized in P sequestration as *Gemmatimonas* [52] and in the acidophilic Planctomycetes *Singulisphaera* [53]. However, among the bacterial *taxa* significantly enriched, many Proteobacteria showed negative correlation being related with many other fungal guilds. This suggests that the association between fungus and bacteria may be more appropriately studied considering species specific interactions. However, the actual data provided in this work do not allow to go in depth of the possible interactions occurring between the two Kingdoms requiring specific experimental settled to solve possible hypotheses.

### Microbial relationships of truffle mycelium in expanding truffle forests

The Spearman correlation rank in Figure 7 illustrates an adverse relationship between the two fungal guilds. Even though the *Tuber* genus is part of the ECM, we observe a distinctly different trend compared to its guild. This could have two explanations, both related to its biological cycle. Firstly, the mycelium’s growth rate appears to be highest in spring, then decreases significantly in summer, and resumes growth in autumn, when it concentrates on producing ascocarps [54]. The sampling for this study was conducted in autumn at random points in each area. The goal was to describe the soil environment independently of the exact point where the white truffle is produced. This might explain a lower presence of the *taxon* in question. However, this could also lead to the hypothesis of different conservation strategies. For example, organisms might move to greater depths in search of underlying water resources and less competition [55]. Another strategy could involve associating with generally non-host plants, assuming endophytic life style as demonstrated for other species of *Tuber*, making it difficult to detect [56]. This hypothesis could also explain the positive correlation identified with endophytes (Cluster 1, Figure 7), which, along with the other associations, needs further investigation. Concerning the specific correlation of *Tuber* mycelium with bacterial *taxa*, the last presents high association with *Ktedonobacter, Zavarzinella* and *Sphingomonas* members of Chloroflexota, Plantomycetota and Pseudomonadota, respectively. For the first two, none interaction with *Tuber spp*. has been described in literature. The former has a wide ecological range, proliferating in both common and extreme environments [57], while the second constitutes a mono specific genus (*Zavarzinella formosa*) recently described as a new species and isolated in peat from boreal environment [58]. Instead *Sphingomonas* genus were found in soil (roots) truffle site and ascomata of *T. aestivum* [59,60]. Those bacteria showed to be able to enhance plant growth and drought resistance through multiple mechanisms, for example has been shown to stimulate the formation of secondary roots [61], which are essential for the formation of new mycorrhizae [62-64]. Closely co-occur with *Tuber melanosporum* Vittad. soil site [65]. Pavić et al in 2011 [66] isolate *Sphingobium* sp. TMG 022C from the ascocarp of *T. magnatum* and demonstrated its ability to perform ammonification and nitrate reduction, solubilise phosphate, hydrolyse lipids and degrade β-glucans and chitin, could thus be involved in mycelium nutrition and ascocarp growth and decomposition.

## Conclusion

In this study, we presented a description of the truffle soil microbiome in a productive forest and the nearby expanding area. The results indicate stable biological communities in the primary forest, which appear to be particularly sensitive to seasonal changes, as evidenced by the differences found between the two sampling years. In the expanding truffle forest, we find bacterial and fungal communities that are not yet well-defined and have greater dynamism in response to environmental changes. Another interesting interaction was observed in the competition between saprophytes and ECMs. The behavior and dynamics of the latter in response to management interventions in the study area show a preference in litter-limited conditions. When we look closely at the dynamics of the *Tuber* genus, we find significant differences in the trophic guilds they belong to. This emphasizes that the interactions between individual *taxa* are influenced by their biology. This suggests that more research is needed to understand the relationships between the specific *taxa* in the environment.

---

**Figure S1:**
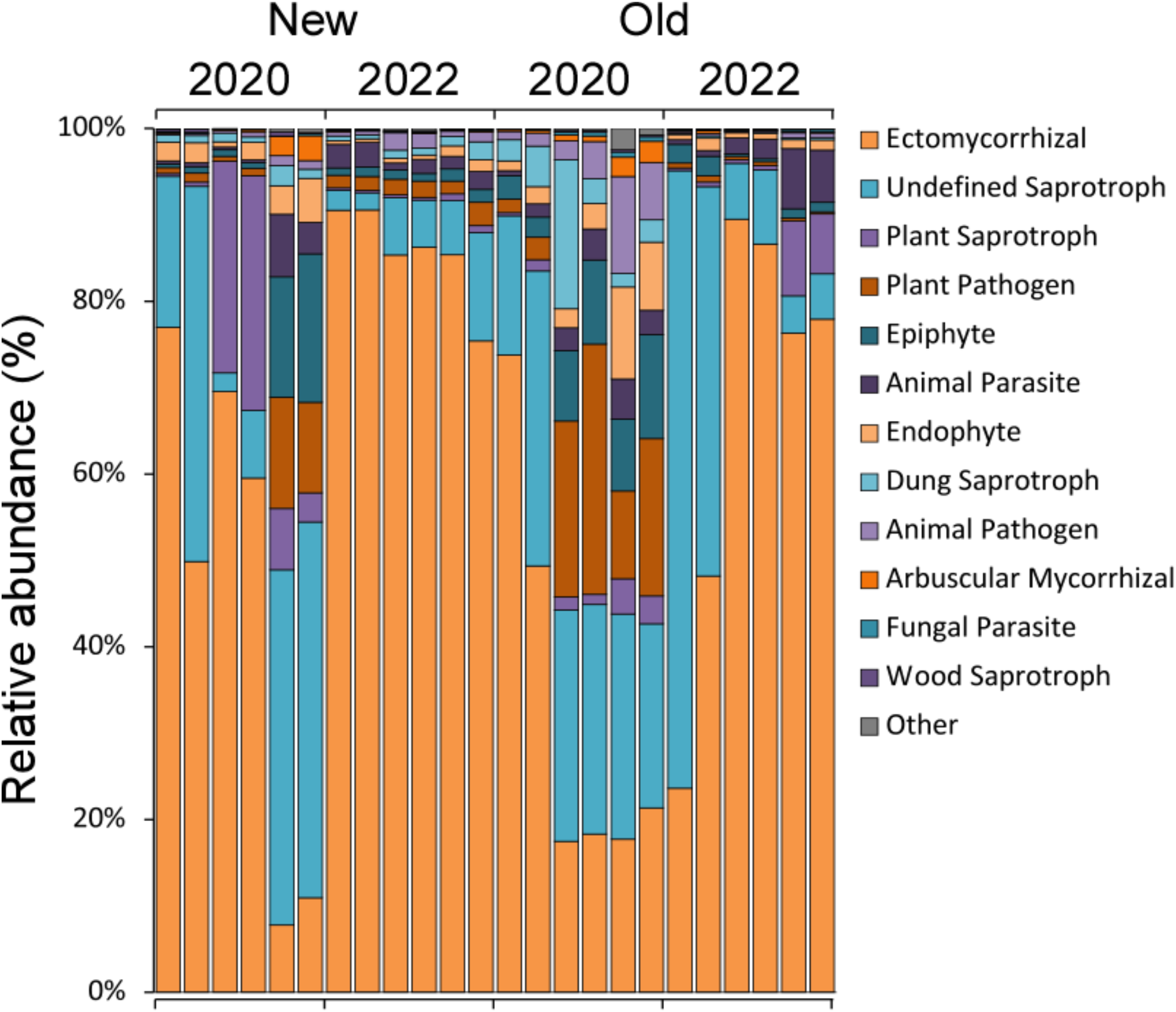
Staked bar plot showing the different patterns in relative abundance of fungal data based on FunGuild trophic strategy associations.

